# Flexible predictive control in human interception under visual occlusion and altered gravity

**DOI:** 10.64898/2026.07.09.737249

**Authors:** Marta Russo, Amine Chaigneau, Giovanni Pezzulo

**Affiliations:** Institute of cognitive sciences and technologies (ISTC), CNR, Via Gian Domenico Romagnosi 18, Roma, 00196, RM, Italy

**Keywords:** internal model, gravity, occlusion, interception, kinematics, motor control, motor behavior

## Abstract

Interception of moving objects requires the nervous system to compensate for sensory delays and uncertainty, yet how behavior is controlled remains debated. Key questions concern whether predictive processes play any role at all and, if so, whether they rely on simple motion extrapolation or incorporate internalized physical priors, such as gravity. Another open question is whether observers adopt a single control strategy or flexibly switch between predictive and reactive control - or between different predictive strategies - depending on task demands. To address these questions, we developed a virtual interception task in which participants intercepted moving targets under systematically varied conditions. We manipulated gravity (**1*g*** vs. **0*g***), visual availability (occluded vs. non-occluded), target velocity, and the initial spatial configuration of the ball and paddle (same vs. opposite side). Results indicate that interception is supported by predictive mechanisms across conditions. Behavioral patterns during occluded **0*g*** trials suggest that participants extrapolate target motion using expectations consistent with gravity. Target velocity, visual occlusion, and task geometry modulated movement strategies, indicating that predictive control is flexibly adapted to task demands. These findings support the view that interception relies on predictive internal models incorporating structured physical priors while revealing flexible, context-dependent adaptations to sensory and task constraints.

## 1 Introduction

Interception of moving objects is a fundamental sensorimotor ability that requires precise visuomotor coordination despite sensorimotor delays, perceptual uncertainty, and motor noise (Wolpert and Flanagan, 2001; Harris and Wolpert, 1998; Shadmehr et al., 2010). To successfully intercept a moving target, the central nervous system (CNS) must guide the effector to the right location at precisely the right time, compensating for delays that make purely reactive control challenging.

Several theories have been proposed to explain how the CNS accomplishes this task. Early accounts emphasized reactive mechanisms, suggesting that actions are triggered directly by optical variables, such as the rate of retinal expansion (Lee, 1976) or time-to-contact information (Tresilian, 1995, 2004, 2005). More recent computational accounts instead propose that the CNS predicts future target states using internal models of object dynamics (Wolpert and Kawato, 1998; Miall and Wolpert, 1996). These ideas have converged into two broad frameworks for interception: reactive (model-free) online control and predictive (model-based) control (Zhao and Warren, 2015; Russo et al., 2025b; Franklin and Wolpert, 2011; Trevinño et al., 2026). In reactive accounts, movements are continuously coupled to incoming sensory information without explicitly estimating the target’s future trajectory (Dessing et al., 2005; Dessing and Craig, 2010; Dessing et al., 2009). In predictive accounts, the CNS compensates for sensory delays by generating internal predictions of future target states that guide ongoing motor commands (Shadmehr and Krakauer, 2008; Jörges and López-Moliner, 2017; Tenenbaum et al., 2011; Friston et al., 2021; Brenner and Smeets, 2018).

Although substantial evidence supports both frameworks, two fundamental questions remain unresolved. First, if interception relies on prediction, what kind of predictive model does the CNS employ? Predictions could arise from relatively simple extrapolation of recent visual motion, or from richer internal models that encode latent physical regularities of the environment. One prominent proposal is that humans exploit prior knowledge of Earth’s gravity to anticipate object motion (McIntyre et al., 2001; Zago et al., 2009; Jörges and López-Moliner, 2017; Aguado and López-Moliner, 2025). This aligns with the broader view that the brain relies on internal forward models tuned to robust environmental invariants to compensate for sensorimotor delays and predict future states (Torricelli et al., 2023). Such gravity-based priors could improve forecasts of target trajectories (Delle Monache et al., 2023), consistent with findings that interception is generally more accurate when targets obey terrestrial gravity (1*g*) than artificial zero-gravity (0*g*) dynamics (Cullen, 2023, 2012). However, whether these physical priors are continuously incorporated into online predictions remains debated (Jörges and Harris, 2026).

Second, does the CNS rely on a single control strategy, or does it flexibly shift between reactive and predictive control depending on task demands? When the target is slow and remains continuously visible, interception may be achieved through reactive strategies that exploit continuously available optic variables (Brouwer et al., 2000, 2002b,a). In contrast, both faster targets and visual occlusion challenge online visual control. In particular, visual occlusion provides a stringent test of predictive control: if movements remain appropriately guided after the target disappears, they must rely on extrapolation of the unseen trajectory rather than continuous visual servo-control. Task geometry may similarly influence the adopted strategy. For example, when the effector and target begin on the same side, interception may be achieved through relatively simple movement heuristics, whereas starting from opposite sides may require anticipating the future interception point.

To address these questions, we developed *StarBall*, a two-dimensional virtual interception task in which participants controlled a paddle and intercepted balls moving under different physical and sensory conditions. We manipulated ball velocity (small, medium, fast), gravitational acceleration (1g vs. 0g), visual availability (occluded vs. not occluded), and the initial spatial configuration of the target and effector (same side vs. opposite sides), while recording full movement trajectories. If interception relies on predictive models incorporating gravity priors, participants should perform better under 1g motion and exhibit systematic biases during 0g occluded trials, attempting to intercept the target near the location predicted under terrestrial gravity. Furthermore, if predictive control operates irrespective of visual availability, movement kinematics should remain qualitatively similar in occluded and non-occluded trials. In contrast, if participants rely primarily on reactive control when visual information is continuously available, distinct kinematic patterns should emerge between visible and occluded conditions. Finally, differences between same-side and opposite-side starting configurations would indicate that interception strategies depend on task geometry rather than reflecting a single invariant control policy.

## 2 Methods

### 2.1 Participants

Fifty participants (25 women, 25 men) were recruited via the Prolific online platform. All were native English speakers, right-handed, and reported normal or corrected-to-normal vision. The median age was 35 years. Participants completed the task on a desktop or laptop computer equipped with a mouse. Additional technical requirements and compatibility details are reported in Appendix. All participants provided informed consent prior to the experiment. The study was conducted in accordance with the Declaration of Helsinki and approved by [anonymized for review]. Participants were unaware of the hypotheses and purpose of the study.

### 2.2 Experimental design and task

Participants performed a visually guided interception task implemented as a two-dimensional (2D) video game (Figure 1), in which they controlled a paddle using a computer mouse to intercept a falling ball. The experiment followed a fully crossed 3*×*2*×*2*×*2 within-subjects factorial design manipulating *velocity* (slow *v*_1_, medium *v*_2_, fast *v*_3_), *gravity* (gravity 1*g*, no-gravity 0*g*), *occlusion* (occluded *O*, non-occluded *NO*), and *ball–paddle starting configuration* (*Same* side, *Opposite* side). The paddle starting side (*P*^*L*^, *P*^*R*^) was counterbalanced across trials but was not part of the factorial design.

**Fig. 1.**
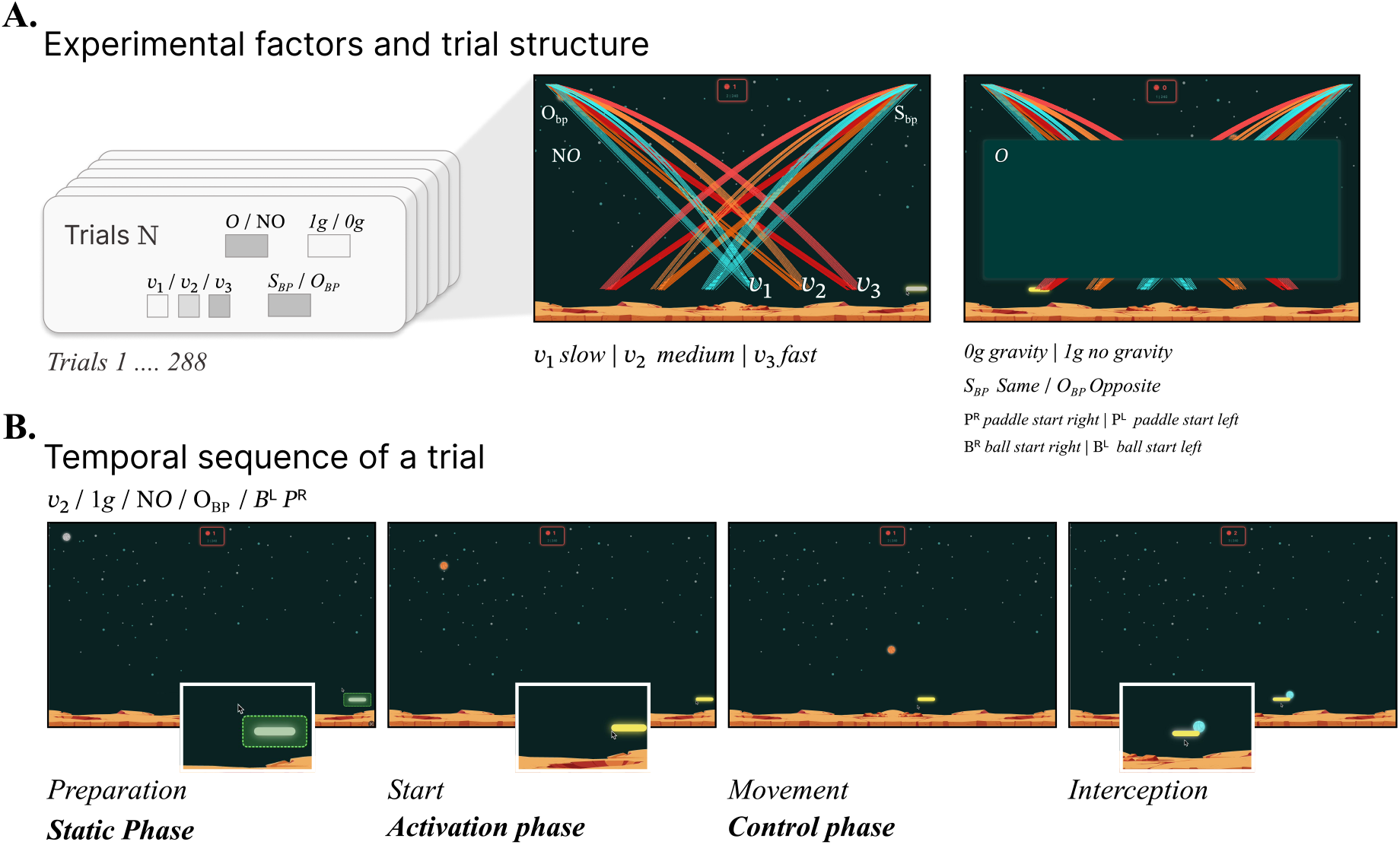
Overview of the experimental task. Panel A shows the main game interface and experimental design. Participants controlled a yellow paddle using the computer mouse to intercept a falling ball within a fixed 2D environment. The experiment followed a fully crossed 3 *×* 2 *×* 2 *×* 2 factorial design manipulating *velocity* (*v*_1_, *v*_2_, *v*_3_), *gravity* (1*g*, 0*g*), *occlusion* (*O, NO*), and *ball starting configuration* (*Same* (*S*_*bp*_), *Opposite* (*O*_*bp*_)). The paddle starting position was counterbalanced across trials (*P*^*L*^, *P*^*R*^). Ball trajectories were defined by initial velocity vectors [*v*_*x*_, *v*_*y*_], resulting in 12 trajectory types distributed across 6 landing areas. Participants completed 288 trials in random order within a single experimental block. Panel B shows the temporal structure of a trial. Each trial comprised three phases: a *static phase*, during which both the ball and paddle remained stationary; an *activation phase*, triggered when the participant’s cursor entered the green activation zone, enabling paddle horizontal control and launching the ball; and a *control phase*, during which the participant moved the paddle horizontally to intercept the falling ball.

Ball trajectories were determined by initial velocity vectors [*v*_*x*_, *v*_*y*_], producing 12 distinct trajectory types distributed across six landing regions (Figure 1A). Participants completed 288 trials in a fully randomized order for each participant within a single session, corresponding to 12 repetitions per condition.

Each trial consisted of three sequential phases (Figure 1B): (1) a *static phase*, during which both ball and paddle remained stationary; (2) an *activation phase*, initiated when the participant moved the cursor into a predefined activation region, which enabled paddle control and launched the ball after a brief randomized delay (uniformly sampled between 0 and 50 ms); and (3) a *control phase*, during which participants attempted to intercept the falling ball by moving the paddle horizontally. The paddle’s vertical position was fixed, such that control was limited to a single horizontal degree of freedom, determined by the cursor’s horizontal position and independent of its vertical movement. Participants received immediate feedback on performance. Successful interceptions caused the ball to stop at the collision point, turn blue, and trigger a success sound; missed interceptions resulted in the ball continuing its trajectory until reaching the bottom of the display, where it turned red and triggered an error sound.

To reduce spatial predictability, the ball’s initial horizontal position varied randomly within a restricted range near the starting side. In occlusion trials (*O*), the ball’s flight trajectory was partially obstructed by a central occluding panel (Figure 1A).

Participants were instructed to catch as many balls as possible. They received a base payment of 6 eand a performance-dependent bonus (0–4 €), which was communicated in advance. The average total payment was approximately 8 *±* 1 €. The experimental session lasted approximately 35 minutes. Prior to the main experiment, participants viewed a short instructional video and completed 12 practice trials (one by experimental condition).

### 2.3 Data processing

Behavioral and trajectory analyses were conducted at the trial level. Catch outcomes were first verified offline using the same spatial criteria as the online collision detection procedure (see Appendix). Trials classified as false negatives (2.93%) were re-coded as successful and truncated at the first catchable frame.

For missed trials, interception error (*error*) was defined as the minimum Euclidean distance between the centers of the ball and paddle over the trial. To characterize paddle–ball coordination, ball and paddle trajectories were resampled onto a uniform temporal grid (0.025s resolution, 40Hz) and smoothed using a Savitzky–Golay filter. For each trial, trajectories were normalized relative to their initial positions and the horizontal crossing point of the ball at paddle height. We then computed a coupling index as the difference between normalized paddle and ball positions over time. A coupling ratio (*AUC*_*ratio*_) was then derived as the normalized balance between positive and negative areas under this difference, providing a bounded summary measure of coordination. Positive values indicate that the paddle leads the ball trajectory on average, whereas negative values indicate that it lags behind.

Cursor trajectories were temporally resampled to 101 normalized time steps using linear interpolation, following standard procedures in cursor movements analysis (Wulff et al., 2025). To facilitate comparison across trials, trajectories were horizontally flipped to a common left-to-right reference frame based on paddle starting side and remapped to a common spatial origin. Vertical control behavior was quantified using the signed maximal deviation (*MD*) of the cursor relative to the paddle reference position (*y* = 107 px). Positive values indicates deviation toward the ball, while negative values indicates deviation below the paddle. Additional preprocessing details and metric definitions are reported in Appendix.

### 2.4 Data analysis methods

Task performance (*success*) was analyzed using generalized linear mixed-effects models (GLMMs) with a logit link, fitted in *R* using the *lme4* package (Bates et al., 2015). The primary model tested the effects of *occlusion, gravity, velocity*, and *starting configuration*, including the *occlusion × gravity* interaction, while accounting for repeated observations within participants via a random intercept. Pairwise contrasts were computed for the *occlusion × gravity* interaction using estimated marginal means (*emmeans*). Participant-level variables (e.g., age, gender) were not included in the main model, as they were not central to the hypotheses, but were examined in supplementary analyses reported in the Appendix.

*Error, RT, MD* and *AUC*_*ratio*_ were analyzed using linear mixed-effects models (LMMs) with the same fixed-effects structure as the primary performance model. To account for individual variability in sensitivity to experimental manipulations, random slopes for *gravity* and *occlusion* were included. Effects involving the *occlusion × gravity* interaction were further examined using pairwise contrasts.

For *MD* the random-effects structure was simplified by removing the random slope for *occlusion* due to convergence constraints. For the AUC_ratio_ (Same) model, the random slope for *gravity* was removed due to a singular fit (*r* = 1.00, *σ*^2^ *<* 0.001). To assess the robustness and generality of the main effects, we conducted additional analyses including interactions with *velocity* and *starting configuration*. Planned contrasts were computed to compare specific condition pairs (e.g., *O* 0*g* vs. *NO* 0*g, O* 1*g* vs. *NO* 1*g*).

Fixed-effect estimates for all models, including odds ratios with 95% confidence and standardized coefficients (*β*^*∗*^) with 95% confidence intervals for all LMMs, are reported in Appendix (Table 1) as well as Random effects variance components (Table 3).

**Table 1.**
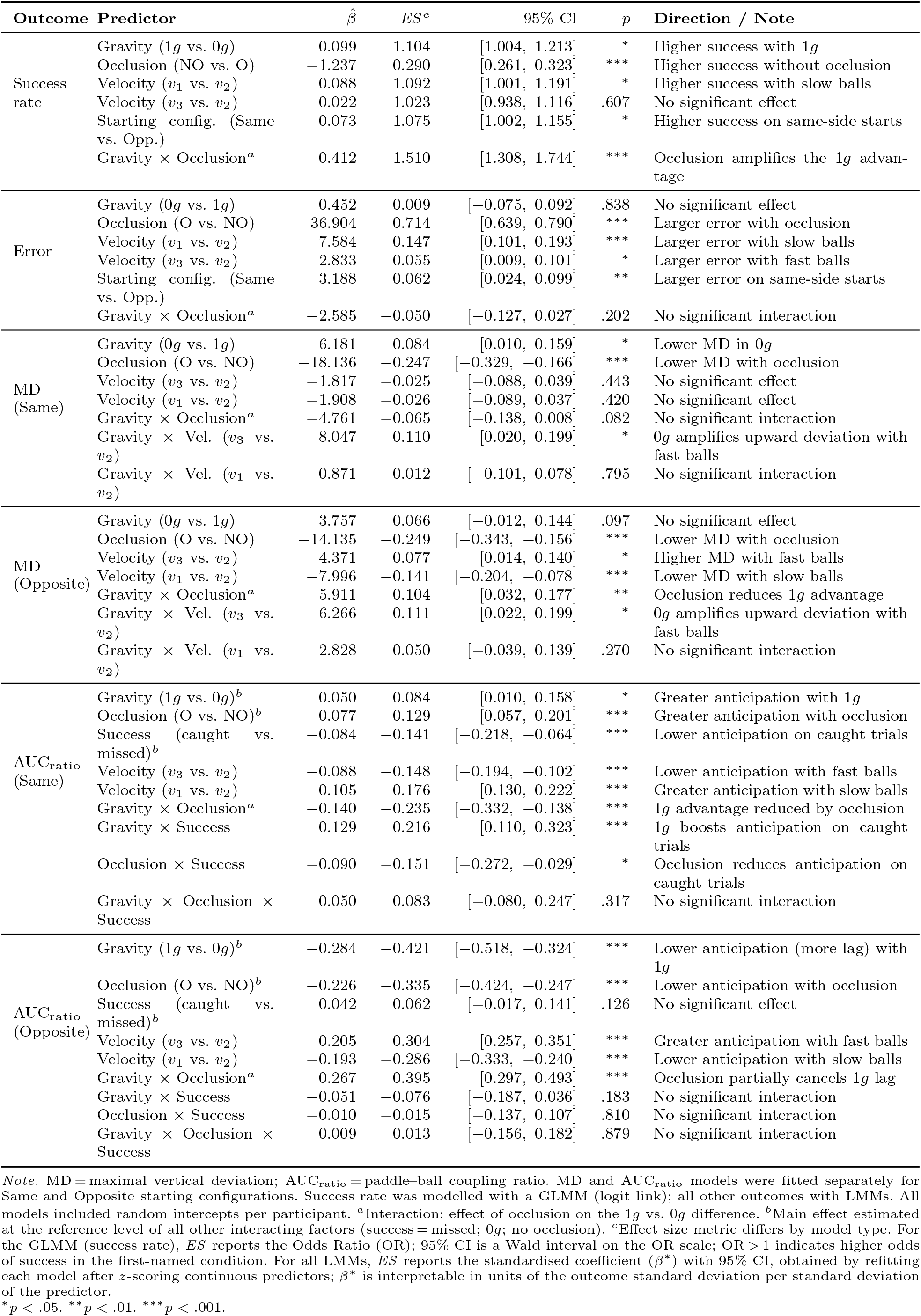
Fixed-effect estimates for all statistical models.

**Table 2.** Pairwise contrasts for AUC_ratio_ models.

| Outcome | Contrast | $\hat{b}$ | $d$ | 95% CI | $p$ | Direction / Note |
| --- | --- | --- | --- | --- | --- | --- |
| $AUC_{ratio}$<br>(Same) —<br>Gravity ×<br>Occlusion<sup>a</sup> | $0g, NO$ vs. $1g, NO$ | −0.115 | −0.238 | [−0.305, −0.170] | *** | Lower anticipation in $0g$ without occlusion |
| | $0g, NO$ vs. $0g, O$ | −0.032 | −0.066 | [−0.151, 0.018] | .308 | No significant difference |
| | $0g, NO$ vs. $1g, O$ | −0.032 | −0.065 | [−0.144, 0.014] | .308 | No significant difference |
| | $1g, NO$ vs. $0g, O$ | 0.083 | 0.172 | [0.087, 0.256] | *** | Higher anticipation in $1g$ without occlusion |
| | $1g, NO$ vs. $1g, O$ | 0.083 | 0.172 | [0.094, 0.251] | *** | Higher anticipation in $1g$ without occlusion |
| | $0g, O$ vs. $1g, O$ | 0.000 | 0.001 | [−0.077, 0.078] | .985 | No significant difference |
| $AUC_{ratio}$<br>(Same) —<br>Success<sup>b</sup> | $0g, NO$: Caught — Missed | −0.082 | −0.170 | [−0.268, −0.072] | *** | Lower anticipation on caught trials |
| | $1g, NO$: Caught — Missed | 0.042 | 0.088 | [−0.009, 0.185] | .070 | No significant difference |
| | $0g, O$: Caught — Missed | −0.174 | −0.360 | [−0.479, −0.241] | *** | Lower anticipation on caught trials; occlusion amplifies effect |
| | $1g, O$: Caught — Missed | 0.003 | 0.006 | [−0.095, 0.108] | .899 | No significant difference |
| $AUC_{ratio}$<br>(Opposite) —<br>Gravity ×<br>Occlusion<sup>a</sup> | $1g, NO$ vs. $0g, NO$ | −0.310 | −0.557 | [−0.657, −0.456] | *** | Lower anticipation (more lag) with $1g$ without occlusion |
| | $0g, O$ vs. $0g, NO$ | −0.231 | −0.415 | [−0.516, −0.315] | *** | Lower anticipation with occlusion in $0g$ |
| | $0g, O$ vs. $1g, NO$ | 0.079 | 0.141 | [0.028, 0.254] | * | Slightly higher anticipation in $0g$ with occlusion vs. $1g$ without |
| | $1g, O$ vs. $0g, NO$ | −0.270 | −0.485 | [−0.620, −0.350] | *** | Lower anticipation with $1g$ and occlusion |
| | $1g, O$ vs. $1g, NO$ | 0.040 | 0.072 | [−0.027, 0.170] | .296 | No significant difference |
| | $1g, O$ vs. $0g, O$ | −0.039 | −0.069 | [−0.178, 0.039] | .296 | No significant difference |

Note. $\hat{b}$ = estimated difference; $d$ = Cohen's $d$ with 95% CI; $p$ = Holm-corrected $p$ -value. <sup>a</sup>Pairwise contrasts decomposing the significant Gravity × Occlusion interaction, marginalised over Success. <sup>b</sup>Contrasts decomposing the significant Gravity × Success and Occlusion × Success interactions; estimated within each Gravity × Occlusion cell. No Success contrasts are reported for *Opposite*, as neither Gravity × Success nor Occlusion × Success reached significance in the omnibus model. \* $p < .05$ . \*\* $p < .01$ . \*\*\* $p < .001$ .

**Table 3.** Random effects variance components for all statistical models.

| Outcome | Groups | Parameter | Variance | SD | Correlations |  |
| --- | --- | --- | --- | --- | --- | --- |
| Success rate | Participant | Intercept | 0.176 | 0.419 |  |  |
| Error | Participant | Intercept | 84.68 | 9.202 |  |  |
|  |  | Gravity | 119.80 | 10.946 | .18 |  |
|  |  | Occlusion | 98.73 | 9.936 | -.22 | -.46 |
|  | Residual |  | 2214.30 | 47.056 |  |  |
| MD<br>(Same) | Participant | Intercept | 2085.37 | 45.666 |  |  |
|  |  | Gravity | 12.68 | 3.561 | -.11 |  |
|  |  | Occlusion | 280.80 | 16.757 | -.38 | .27 |
|  | Residual |  | 3360.13 | 57.967 |  |  |
| MD<br>(Opposite) | Participant | Intercept | 1533.07 | 39.154 |  |  |
|  |  | Gravity | 35.49 | 5.957 | -.93 |  |
|  |  | Occlusion | 254.90 | 15.966 | -.49 | .40 |
|  | Residual |  | 1971.81 | 44.405 |  |  |
| AUC<sub>ratio</sub><br>(Same) | Participant | Intercept | 0.112 | 0.335 |  |  |
|  |  | Occlusion | 0.004 | 0.064 | .04 |  |
|  | Residual |  | 0.233 | 0.482 |  |  |
| AUC<sub>ratio</sub><br>(Opposite) | Participant | Intercept | 0.081 | 0.284 |  |  |
|  |  | Gravity | 0.021 | 0.147 | .11 |  |
|  |  | Occlusion | 0.018 | 0.133 | -.02 | .29 |
|  | Residual |  | 0.310 | 0.557 |  |  |

*Note.* SD = standard deviation of the random effect distribution. The success rate model (GLMM) included a random intercept only; no residual variance is reported for GLMMs as it is fixed on the latent scale. All LMMs included random slopes for gravity and occlusion per participant, with the exception of MD (Same) and AUC (Same), for which the random slope for occlusion were retained following singular fit. Number of observations and participants: Success rate: $N = 14,397$ , $n = 50$ ; Error: $N = 9,062$ , $n = 50$ ; MD (Same): $N = 7,199$ , $n = 50$ ; MD (Opposite): $N = 7,198$ , $n = 50$ ; AUC<sub>ratio</sub> (Same): $N = 7,199$ , $n = 50$ ; AUC<sub>ratio</sub> (Opposite): $N = 7,198$ , $n = 50$ .

## 3 Results

### 3.1 Performance and errors

Overall, the task resulted in difficulty as participants successfully hit about 50% of the trials. The presence of gravity in the ball trajectory significantly modulated their performance, with higher score when ball trajectories were affected by gravity (*β* = 0.09861, *p <* 0.05). In addition, participants caught more balls when the trajectory was not occluded (*β* = *−*1.23686, *p <* 0.001). However, results of the LMM indicated that between gravity and occlusion factors there was a significant interaction, suggesting that the presence of occlusion increased the difference of caught balls between 1*g* and 0*g* conditions (*β* = 0.41223, *p <* 0.001; see Figure 2A).

**Fig. 2.**
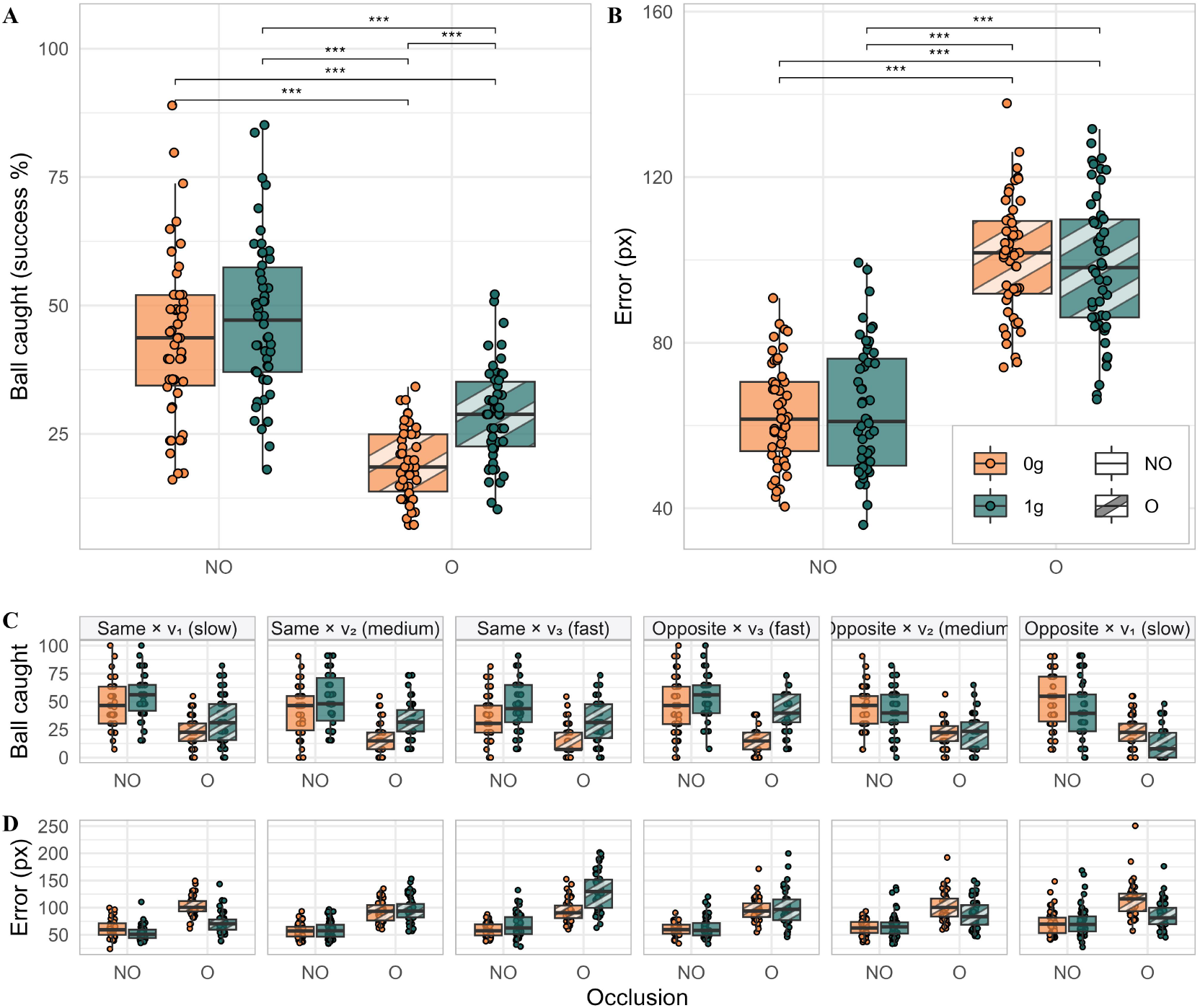
Behavioral performance across experimental conditions. Panel A shows the percentage of successful interceptions (ball caught) as a function of *gravity* (0*g*, 1*g*) and *occlusion* (occluded: *O*, non-occluded: *NO*). Boxplots represent the distribution of participant-level data, with individual data points overlaid. The 0*g* condition is shown in orange and the gravity 1*g* condition in green; *O* and *NO* conditions are distinguished by pattern style. Panel B shows the interception error (*error* in px) for missed trials only, defined as the minimal Euclidean distance between the center of the ball and the center of the paddle. Boxplots and overlaid points follow the same conventions as in Panel A. Asterisks indicate significant pairwise comparisons following the gravity *×* occlusion interaction (Holm-corrected): * *p <* .05, ** *p <* .01, *** *p <* .001. Panels C and D show the same dependent measures (*success* rate and *error*, respectively) as a function of *starting configuration* (*Same, Opposite*) and *velocity* (*v*_1_, *v*_2_, *v*_3_). These panels are provided for descriptive purposes only; the corresponding interactions were not included in the statistical models, and no inferential statistics are reported.

Error analysis on missed trials revealed a partially convergent pattern. Spatial error was significantly greater for occluded (*O*) than for non-occluded (*NO*) trajectories (*β* = 36.9041, *p <* 0.001), whereas neither the main effect of gravity (*β* = 0.4517, *p* = 0.83792) nor its interaction with occlusion (*β* = *−*2.5848, *p* = 0.20238) reached statistical significance. Figure 2B shows error distributions across the four experimental conditions. Figure 2C and 2D further shows the effect of velocity and ball starting configuration on spatial error and success.

Taken together, these results show that interception became more challenging under both visual occlusion and 0*g* motion, with the combination of these conditions producing the largest performance costs. This pattern suggests that participants relied on prior expectations of natural (1*g*) gravity to guide paddle movements, particularly when the task was most demanding.

### 3.2 Paddle kinematics, horizontal movements

In order to evaluate the horizontal movement of the paddle in relation to ball motion, we defined the coupling metric as the difference between the paddle and ball horizontal position at the same time. Due to the different configuration of *Same* and *Opposite* experimental conditions, we divided the data set in these two subsets and examined them separately.

Figure 3 shows the results for the *Same* condition, i.e. when both ball and paddle started from the same side. The ratio of the AUC above (and below) the zero-line indicates how much the paddle anticipated (followed) the ball. In Figure 3A the ratio of the coupling AUC is shown for caught and missed balls. It shows that across all conditions, participants anticipate the ball rather than just following it. This metric was higher with the occluded 1*g* ball trajectory, as indicated by the significant results of the LMM for the occlusion factor (*β* = 0.077, *p <* 0.001), the gravity factor (*β* = 0.050, *p <* 0.05) and their interaction (*β* = *−*0.140, *p <* 0.001). In addition, the anticipation ratio was different for caught and missed balls (*β* = *−*0.084, *p <* 0.001), and interacted significantly with gravity (*β* = 0.129, *p <* 0.001), suggesting that participants anticipated more in 1*g* and caught more balls. Instead, occlusion reduced anticipatory behavior on caught balls (*β* = *−*0.090, *p <* 0.05).

**Fig. 3.**
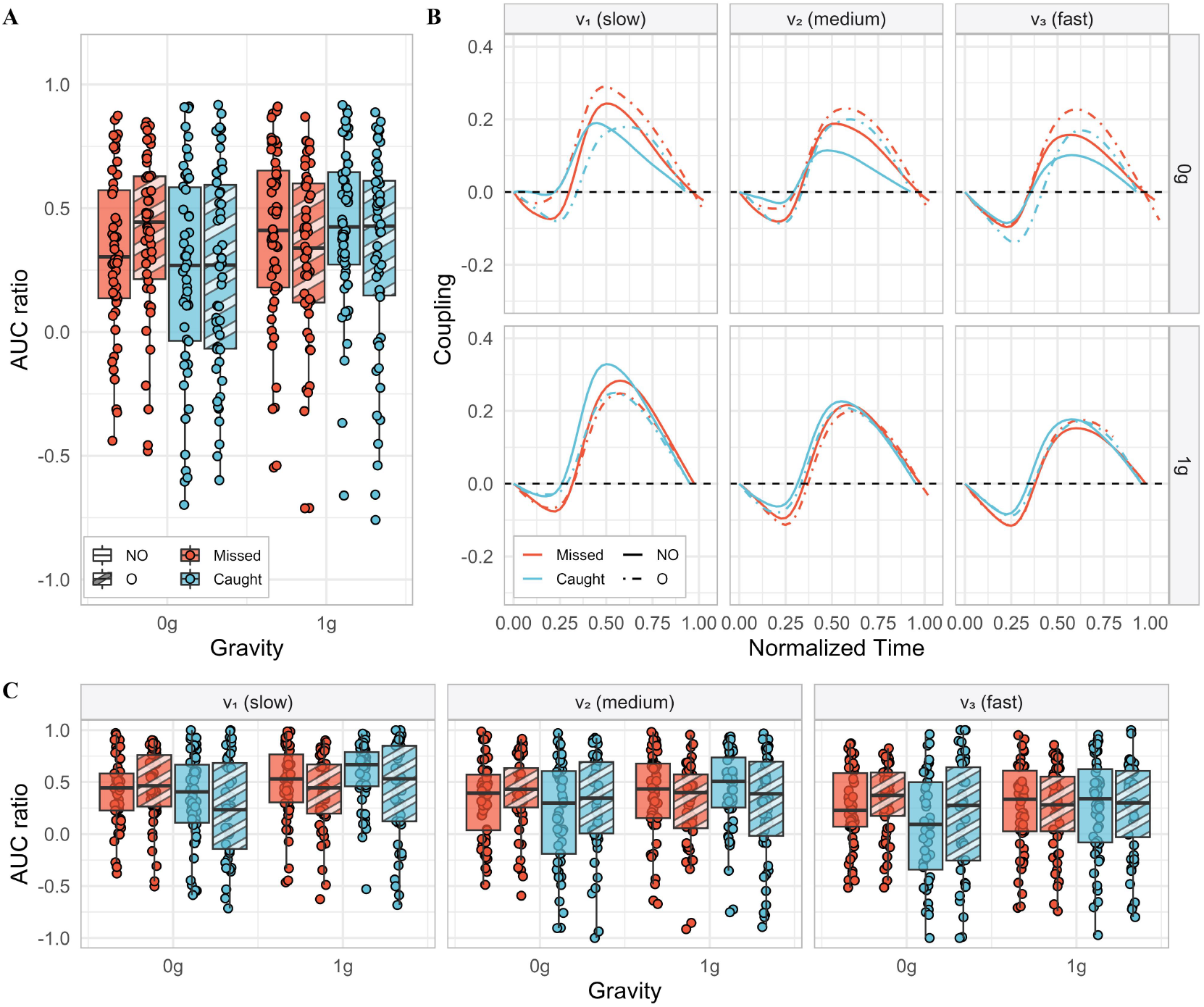
Paddle–ball coupling dynamics in the *Same* starting configuration. Panel A shows the coupling ratio (*AUC*_*ratio*_) as a function of *gravity* (0*g*, 1*g*), *occlusion* (*O, NO*), and *success* (*Caught, Missed*). Boxplots represent participant-level distributions with individual observations overlaid. Positive values indicate that the paddle led the ball trajectory on average, whereas negative values indicate that it lagged behind. Panel B shows the time-resolved coupling index, defined as the difference between normalized paddle and ball positions, across normalized trial time (0–1), as a function of *velocity* (*v*_1_ slow, *v*_2_ medium, *v*_3_ fast; columns) and *gravity* (rows). Lines represent the mean across participants. The horizontal dashed line indicates zero coupling (perfect alignment between paddle and ball). Conditions are defined by *occlusion* (*NO*, solid lines; *O*, dot-dashed lines) and *success* (*Caught*, blue; *Missed*, red). Panel C shows the coupling ratio (*AUC*_*ratio*_) as a function of *gravity, occlusion*, and *success*, separately for each *velocity*. Boxplots represent participant-level distributions with individual observations overlaid. This panel is provided for descriptive purposes only, as the corresponding interaction was not included in the statistical model.

Temporal profiles of the coupling metric are displayed in Figure 3B to illustrate how participants’ behavior changed with time. This figure illustrates that after a short initial period in which it lags behind the ball, the paddle moves ahead of the ball for most of the trial. Furthermore, a noteworthy difference between the 0*g* and the 1*g* conditions emerge: in the former, but not in the latter, the temporal profiles during occluded and not occluded trials are markedly different. Interestingly, the post-hoc comparisons show that the coupling in 0*g* with the *O* for the missed trials was different from the coupling in the 0*g* caught trials (*O*: *β* = 0.174, *p <* 0.001; *NO*: *β* = 0.160, *p <* 0.001) and similar to the coupling in the 1*g −NO* caught trials (*β* = *−*0.017, *p* = 0.99). This result suggests that with *Occlusion* in 0*g* participants engaged in the same prediction for the condition 1*g*. Finally, the AUC ratio was modulated by velocity of the ball as described in Figure 3C.

Participants appear to anticipate the ball also in the case in which the paddle started from the opposite side of the ball (see Figure 4A). In this condition, a strong effect of velocity was observed, although the trend was opposite to that of the previous condition: participants anticipated more when the ball moved faster (*v*_3_ vs *v*_2_ *β* = 0.205, *p <* 0.001; *v*_1_ vs *v*_2_ *β* = *−*0.193, *p <* 0.001). A possible explanation is that at the highest ball velocity, the distance covered by the paddle was smaller, hence (after normalization) it could appear to anticipate more the ball. Gravity had an opposite effect as well, increasing anticipatory behavior in 0*g* (*β* = *−*0.284, *p <* 0.001) and so did *Occlusion* which elicited less anticipation (*β* = *−*0.226, *p <* 0.001) especially in the 0*g* condition (*β* = 0.267, *p <* 0.001). Notably, the *v*_3_(*slow*) condition with 1*g* acceleration is the only condition in which there is almost no anticipation, with the paddle moving at the same speed of the ball, as shown in the bottom left panel of Figure 4B. Finally, the AUC ratio was modulated by velocity of the ball as described in Figure 3C.

**Fig. 4.**
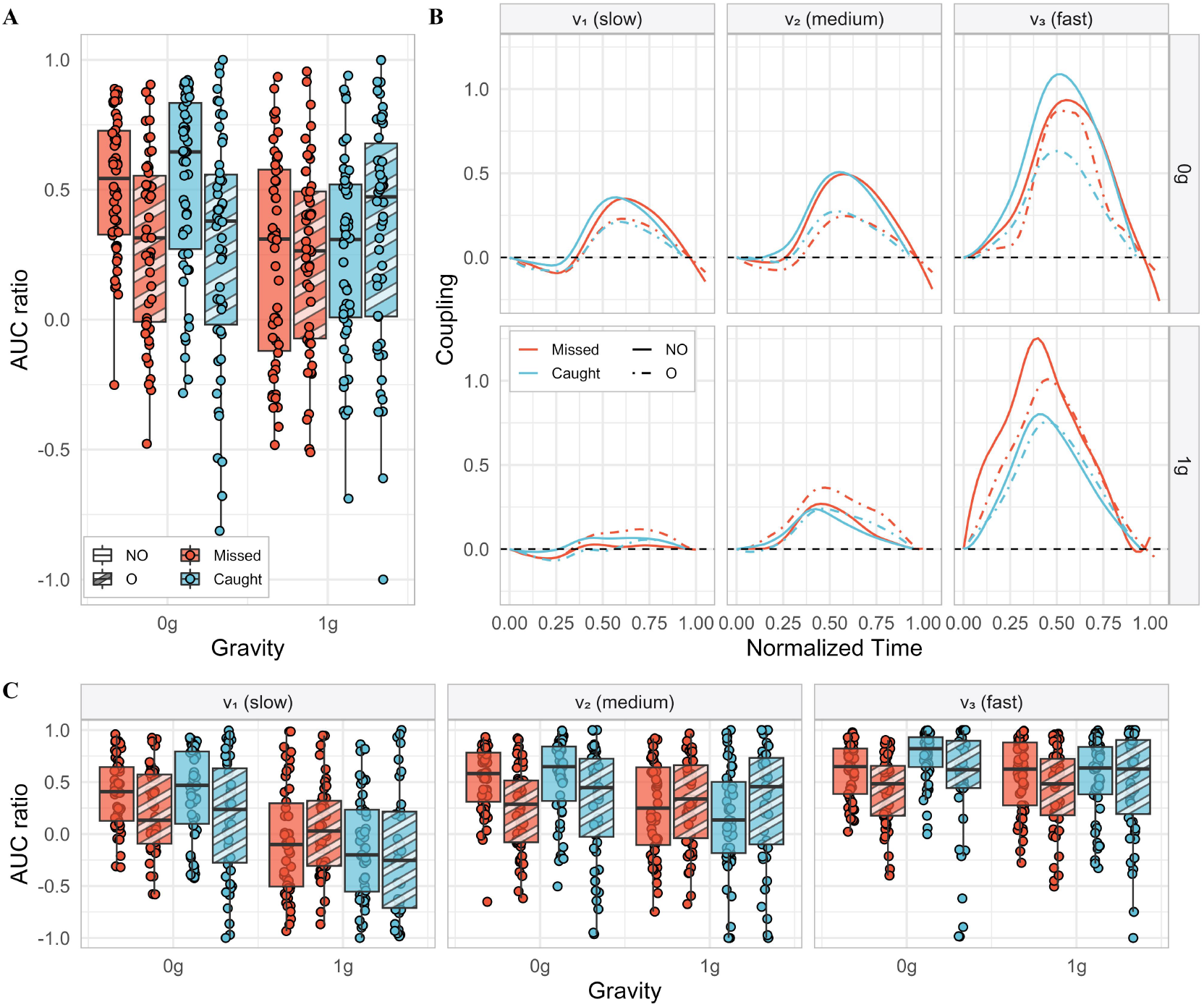
Paddle–ball coupling dynamics in the *Opposite* starting configuration. Panel A shows the coupling ratio (*AUC*_*ratio*_) as a function of *gravity* (0*g*, 1*g*), *occlusion* (*O, NO*), and *success* (*Caught, Missed*). Boxplots represent participant-level distributions with individual observations overlaid. Positive values indicate that the paddle led the ball trajectory on average, whereas negative values indicate that it lagged behind. Panel B shows the time-resolved coupling index, defined as the difference between normalized paddle and ball positions, across normalized trial time (0–1), as a function of *velocity* (*v*_1_ slow, *v*_2_ medium, *v*_3_ fast; columns) and *gravity* (rows). Lines represent the mean across participants. The horizontal dashed line indicates zero coupling (perfect alignment between paddle and ball). Conditions are defined by *occlusion* (*NO*, solid lines; *O*, dot-dashed lines) and *success* (*Caught*, blue; *Missed*, red). Panel C shows the coupling ratio (*AUC*_*ratio*_) as a function of *gravity, occlusion*, and *success*, separately for each *velocity*. Boxplots represent participant-level distributions with individual observations overlaid. This panel is provided for descriptive purposes only, as the corresponding interaction was not included in the statistical model.

Taken together, these results indicate that participants relied predominantly on predictive control across conditions, target velocities, and spatial configurations, with one notable exception (when the ball and the paddle started from opposite sides and the ball moved slowly in 1*g*). When the paddle and ball started on the same side, participants extrapolated occluded trajectories as though the ball obeyed terrestrial gravity (1*g*), even when it did not. Although the same- and opposite-side configurations elicited partially different control strategies, both preserved key signatures of predictive control.

### 3.3 Vertical movements of the cursor

While the paddle could move only horizontally, participants controlled a cursor that also had vertical freedom. Although vertical cursor movement was irrelevant for task success, it was systematically modulated by the experimental conditions. Figure 5 shows the vertical cursor position, with zero corresponding to the paddle’s horizontal position. Maximum vertical deviations are reported in Figure 5A, while average vertical displacement over time is shown in Figure 5B for each experimental condition. Notably, the cursor often deviated above or below the paddle’s horizontal axis.

**Fig. 5.**
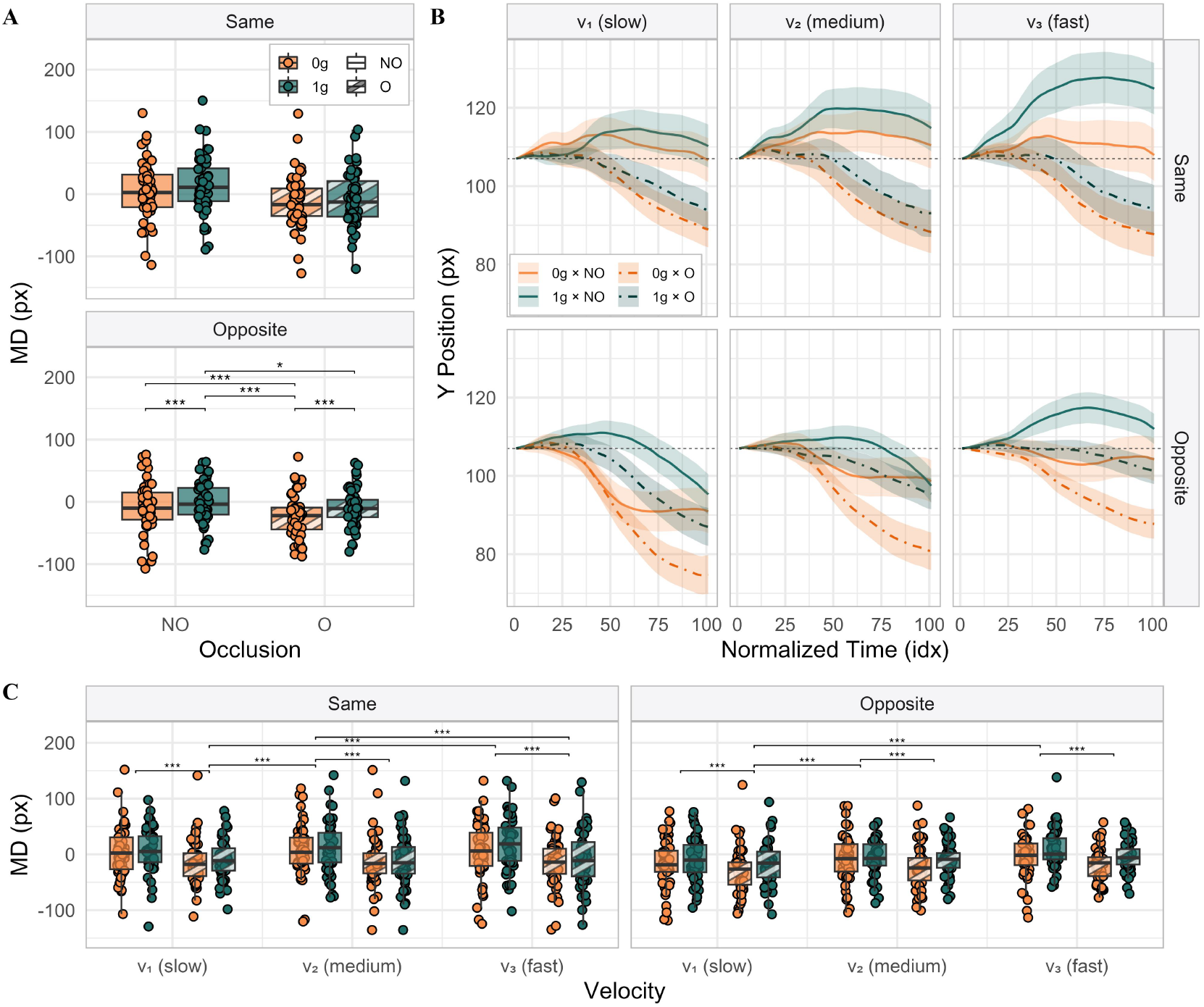
Cursor deviation and trajectory dynamics as a function of task conditions. Panel A shows the signed maximal deviation (MD; in pixels) from the reference paddle position (*y* = 107 px) as a function of *occlusion* (*O, NO*) and *gravity* (0*g*, 1*g*), shown separately for each *starting configuration* (*Opposite, Same*). Boxplots represent the distribution of participant-level data, with individual data points overlaid. Asterisks in the *Opposite* configuration denote significant pairwise contrasts derived from the Gravity *×* Occlusion interaction; no pairwise comparisons were conducted for the *Same* configuration, as the corresponding interaction did not reach significance (*p* = .082). Panel B shows the time-resolved cursor trajectories showing the average vertical position (in pixels) across normalized trial time (0–100), as a function of *velocity* (*v*_3_ fast, *v*_2_ medium, *v*_1_ slow; columns) and *starting configuration* (rows). Lines represent the mean trajectory across participants, and shaded areas denote variability (± SEM). The horizontal dashed line indicates the reference paddle position. Panel C shows the MD as a function of *velocity, gravity*, and *occlusion*, shown separately for each *starting configuration*. Boxplots represent participant-level distributions with individual observations overlaid. Conditions are defined by *gravity* (0*g*, orange; 1*g*, green) and *occlusion* (*NO*, solid lines; *O*, dot-dashed lines). Asterisks indicate significant pairwise contrasts (Holm-corrected) derived from linear mixed-effects models including *gravity × occlusion* and *velocity × occlusion* interactions. Analyses were conducted separately for each starting configuration (*Opposite, Same*), and pairwise comparisons were performed only for significant interactions: * *p <* .05, ** *p <* .01, *** *p <* .001.

As *Same* and *Opposite* side engaged different motor strategies, we analyzed the two conditions with two different LMMs. When ball and paddle started from the *Same* starting configuration, the maximum deviation of the cursor from the paddle movement line was lower in 0*g* (*β* = 6.1807, *p <* 0.05) and with the occlusion (*β* = *−*18.1363, *p <* 0.001), although the interaction between those two factors was not significant (*β* = *−*4.7614, *p* = 0.0815).

The LMM carried out on the *Opposite* data set revealed that maximum deviation in medium velocity condition was significantly different from slow (*β* = *−*7.996, *p <* 0.001) and fast velocities (*β* = 4.371, *p <* 0.05), with slow velocity eliciting the lowest excursions of the cursor. Similarly to the *Same* condition participants moved below the paddle movement line when the trajectory was occluded (*β* = *−*14.135, *p <* 0.001). However, in the *Opposite* scenario gravity was not a significant factor (*β* = 3.757, *p* = 0.09674). Pairwise comparisons for both *Same* and *Opposite* conditions are shown in Figure 5C.

Taken together, these results suggest that participants used vertical cursor movement as an additional degree of freedom that, although task-irrelevant, was systematically modulated by task demands. A possible interpretation of the increased downward movement during occluded trials is that participants attempted to compensate for uncertainty by “buying time,” effectively trying to intercept the ball below the cursor trajectory. However, such a strategy could not improve performance in the present task, where interception required alignment with the target rather than vertical offset.

## 4 Discussion

In this study, we investigated whether people rely on predictive mechanisms to intercept moving objects and, if so, what form these predictions take. Specifically, we asked whether interception relies on simple motion extrapolation or on richer internal models that encode physical regularities such as terrestrial gravity, and whether the adopted control strategy remains invariant or flexibly adapts to task demands. Using the *StarBall* paradigm, in which participants controlled a paddle to intercept falling balls, we manipulated target velocity (fast, medium, slow), gravitational dynamics (1*g* vs. 0*g*), visual availability (occluded vs. non-occluded), and the initial spatial configuration of the ball and paddle (same vs. opposite side). Three main findings emerged.

First, our results provide converging evidence that interception is predominantly supported by predictive control rather than purely reactive online guidance (Friston, 2011; Adams et al., 2013; Parr et al., 2021). Although both visual occlusion and the absence of gravity (0*g*) impaired interception performance, kinematic analyses revealed anticipatory behavior across nearly all conditions. Following an initial tracking phase, participants typically moved the paddle ahead of the ball for most of its trajectory, indicating that they were aiming toward a future interception point rather than continuously following the target (Diaz et al., 2013; Hayhoe et al., 2012; Land and McLeod, 2000). Consistent with this interpretation, the paddle–ball coupling ratio was higher under natural gravity (1*g*), suggesting that target trajectories obeying familiar physical laws facilitated anticipatory control. Computationally, these movement patterns resemble strategies in which the controller predicts the future interception location rather than continuously tracking the target’s instantaneous position (Russo et al., 2025a). A notable exception occurred when the paddle and ball started on opposite sides and the ball followed a 1*g* trajectory, where paddle–ball coupling approached unity and anticipatory behavior was largely absent. This finding suggests that, under relatively simple kinematic conditions, continuous visual servo-control may suffice for successful interception.

Second, our findings suggest that predictive control incorporates internalized expectations about terrestrial gravity. Performance deteriorated under both visual occlusion and 0*g* motion, with the largest costs occurring when these manipulations were combined. Importantly, participants’ behavior during occluded 0*g* trials closely resembled that observed under 1*g*, indicating that they behaved as though the hidden target continued to obey Earth’s gravity. Because this expectation was inconsistent with the actual target dynamics, it produced systematic and predictable interception errors. These findings are consistent with previous work proposing that interception relies on internal models incorporating stable physical regularities, particularly when sensory information is uncertain or unavailable (La Scaleia et al., 2015; Russo et al., 2017; Delle Monache et al., 2023; Neupärtl et al., 2021, 2020). More generally, they support the view that uncertain sensory information is integrated with strong experience-based priors to guide prediction, a principle that has also been proposed to underlie expert performance in dynamic sports such as tennis and baseball (Maselli et al., 2023; Gordon et al., 2021; Brantley and Körding, 2024; Beck et al., 2025).

Third, our results indicate that predictive control is flexible rather than stereotyped. Although predictive signatures were preserved across nearly all experimental conditions, the specific control strategy adapted to task demands. The same- and opposite-side configurations elicited partially different movement kinematics while maintaining anticipatory behavior, suggesting that participants adjusted how prediction was implemented rather than switching wholesale between predictive and reactive control (Iodice et al., 2015). Furthermore, the differences observed between same-side and opposite-side configurations indicate that the adopted predictive control is flexibly adapted to the spatial geometry of the task, consistent with recent findings that complex motor behaviors rely on embodied planning tailored to specific environmental constraints (Maselli et al., 2025). Likewise, under the most demanding conditions, particularly during visual occlusion, participants systematically deviated the cursor well below the required horizontal interception line, despite this movement having no effect on the one-dimensional paddle trajectory. Although speculative, this behavior may reflect an attempt to “buy time” by aiming for a lower interception point, revealing that participants exploited an otherwise task-irrelevant degree of freedom as uncertainty increased. Indeed, structured kinematic patterns are known to carry rich information revealing hidden cognitive states and predictive plans (Becchio et al., 2024).

Several limitations should be acknowledged. First, the experiment was conducted remotely via an online platform (Prolific), introducing unavoidable variability in participants’ hardware and testing environments. Second, the task involved a one-dimensional mouse-controlled paddle and therefore lacked several features of natural interception, including stereoscopic depth, haptic feedback, and unconstrained multi-joint arm movements (Beck et al., 2025). Consequently, caution is warranted when generalizing these findings to naturalistic catching.

These limitations also motivate several directions for future work. Laboratory implementations combining eye tracking with robotic manipulanda could provide higher-resolution measurements of eye-hand coordination (Cesqui et al., 2015, 2012, 2016) while allowing precise mechanical perturbations to probe predictive control. Such studies could also help disentangle the respective contributions of visual and vestibular signals to gravity-based prediction. Beyond behavioral experiments, identifying the neural and computational mechanisms supporting flexible predictive interception remains an important challenge. Previous work has implicated regions such as the dorsal anterior cingulate cortex (dACC) and posterior parietal cortex (PPC) in representing future target locations during dynamic behavior (Yoo et al., 2021), while recent computational studies have shown that predictive tracking naturally emerges in recurrent neural networks trained on analogous spatial interception tasks (Redman et al., 2026). In addition, recent computational theories suggest that the brain dynamically integrates incoming sensory and task information to flexibly update and mix sensorimotor control policies in real time (Christopoulos and Schrater, 2015; Chiappa et al., 2024; Friston, 2011; Priorelli et al., 2023). Extending these approaches to the richer range of manipulations introduced here—including gravity, visual occlusion, target velocity, and spatial configuration—may provide a more comprehensive account of the neural and computational principles underlying predictive interception.

## Acknowledgments

We would like to thank Prof. Constantin Rothkopf for his valuable feedback on an early version of the manuscript.

## Materials and Data availability

The datasets and scripts used in the current study are available in the Open Science Framework repository, [anonymous repository](https://osf.io/h746p/overview?view_only=1d93321e437446e8af05fe608450aefd). All materials can be found on GitHub upon request [details anonymized for review].

## Author contributions

Conceptualization: MR, GP

Data curation: MR, AC

Formal analysis: MR, AC

Funding acquisition: GP

Investigation (experiments): AC

Methodology: GP, MR, AC

Software: AC

Supervision: GP, MR

Validation: GP, MR

Visualization: AC

Writing (original draft): MR, AC

Writing (review & editing): MR, AC, GP

## Appendix A: Technical and Implementation Details

### Hardware and software environment

Participants were required to complete the experiment on a computer equipped with a mouse and a display resolution between 1440*×*900 and 2880*×*1800 pixels. The task was run in the Google Chrome browser (version 138.0.0.0 or later).

### Display geometry and object parameters

The experimental task was implemented in Next.js (https://nextjs.org/) and rendered at a fixed resolution of 1440*×*900 pixels across participants. The ball measured 30*×*30 pixels and the paddle 70*×*14 pixels, and both object sizes remained constant across trials.

The ball’s initial vertical position was fixed at 40 pixels from the top of the display. Its initial horizontal position varied pseudo-randomly in discrete 5-pixel increments within a 40-pixel window adjacent to the starting side (left starts: *x ∈ {*0, 5, …, 40*}*; right starts: *x ∈ {*1370, 1375, …, 1410*}*). The paddle’s vertical position was fixed at 100 pixels from the bottom of the display (*y* = 800 pixels from the top), and its horizontal starting position was initialized at either 10 pixels from the left edge or 1360 pixels from the left edge (i.e., 70 pixels from the right edge), counterbalanced across trials.

In occlusion trials, a rectangular occluding panel (513 pixels tall, 1320 pixels wide) spanned the full width of the display leaving 60-pixel lateral margins on each side. The panel was anchored 153 pixels above the bottom edge, so that it extended from 153 to 666 pixels from the bottom of the display (i.e., from *y* = 234 to *y* = 747 pixels measured from the top).

### Game parameters

The experiment used a fully crossed factorial design. Three main parameters were varied: *gravity, velocity*, and *occlusion*.

- *Gravity* took one of two values: *g ∈ {*0, 981*}* pixels/s^2^.
- *Initial velocity* was specified as a two-dimensional vector [*v*_*x*_, *v*_*y*_] (pixels/s). The horizontal component *v*_*x*_ took one of three values corresponding to three speed classes:

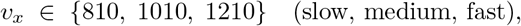

and the vertical component *v*_*y*_ took one of two values:

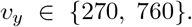

All combinations of *v*_*x*_ and *v*_*y*_ were used, yielding six distinct velocity magnitudes. At runtime, the sign of the horizontal component was reversed for right-side starts (*v*_*x*_ *→ −*|*v*_*x*_|), producing 12 distinct trajectory directions in total.
- *Occlusion condition* was binary (Occlusion vs. No occlusion).
- *Starting configuration* (Ball and paddle starting sides; left vs. right) were also crossed as independent factors.

Each unique combination of the five factors (2 gravity *×* 3 speed *×* 2 vertical velocity *×* 2 ball side *×* 2 occlusion) was repeated 6 times, yielding 288 trials in total (2 *×* 3 *×* 2 *×* 2 *×* 2 *×* 6 = 288). Initial horizontal positions were jittered pseudo-randomly within the 9-step grid described above, ensuring that no two repetitions of the same condition shared the same starting *x* coordinate.

### Rendering pipeline and collision detection

The task was rendered using a frame-synchronized game loop. Mouse cursor positions were sampled using an event-driven procedure and time-stamped, allowing temporal alignment between paddle motion, ball trajectory, and collision events.

Collisions between the falling ball and the participant-controlled paddle were computed in real time using an axis-aligned bounding-box (AABB) method. A collision was registered when the rectangular regions of the ball and paddle overlapped on both axes and was considered valid only if (i) the horizontal overlap exceeded half of the ball’s width and (ii) the lower edge of the ball intersected the vertical extent of the paddle (Figure 1A). These criteria ensured that interceptions were registered only when the ball was contacted from above rather than from below.

### Offline validation of collision detection

All recorded trajectories were re-evaluated offline using the same spatial overlap criteria applied frame by frame. This verification identified potential classification errors: false positives (FP), in which a trial was marked as successful without physical overlap between the ball and paddle, and false negatives (FN), in which a trial was marked as unsuccessful despite a valid overlap during the trajectory. The offline analysis revealed few false positives (2.93%) and a small proportion of false negatives (0.118%). False-negative trials were reclassified as successful, and their trajectories were truncated at the first frame exhibiting valid contact.

**Fig. 1.**
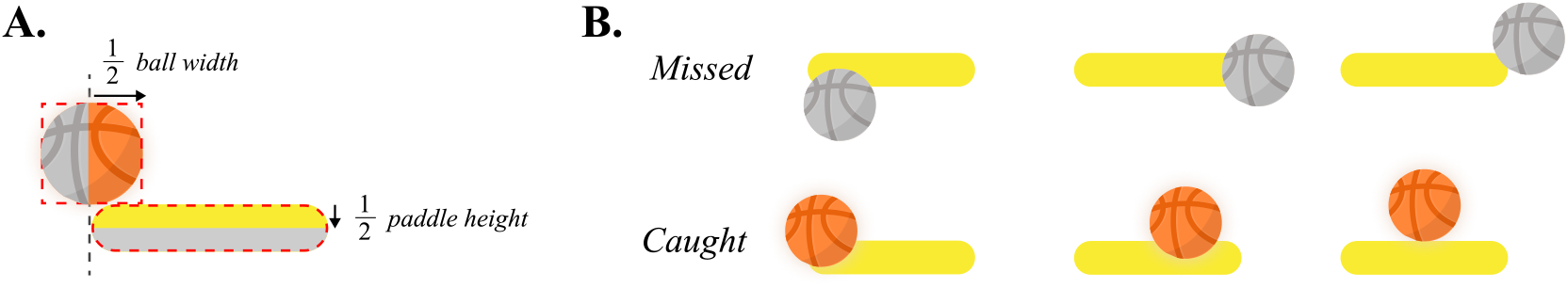
Illustration of collision detection criteria. Panel A depicts the axis-aligned bounding-box (AABB) criteria used to register successful interceptions. A catch was registered when at least half of the ball’s width overlapped horizontally with the paddle and the lower edge of the ball intersected the vertical extent of the paddle, ensuring that the ball could not be caught from below. Panel B provides example outcomes classified as *Missed* (top) or *Caught* (bottom).

## Appendix B: Data processing

### Interception error

For each trial, ball and paddle center coordinates were computed from their rectangular bounding boxes. Interception error was defined as the minimum Euclidean distance between the centers of the ball and paddle over the trial.

### Cursor trajectory resampling and spatial normalization

Cursor trajectories were analyzed from the recorded mouse positions after joining trial-level metadata. For trajectory-based analyses, each trial was resampled to 101 normalized time steps using linear interpolation. To express all trajectories in a common spatial frame, horizontal coordinates were flipped for trials in which the paddle started on the right side, such that movements were represented in a common left-to-right orientation. Trajectories were then remapped to a common origin by subtracting the initial cursor position (20, 107) and translating the result to a fixed paddle-centered reference frame.

### Deviation metric

Vertical deviation was computed at each timestep relative to a fixed reference line corresponding to paddle *y* position (*d*_*i*_ = *y*_*i*_ *− y*_*ref*_). We then computed the signed maximal deviation (*MD*) that was defined as the signed deviation at the time point of maximal absolute deviation 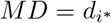, *i*^*∗*^ = arg max_*i*_ |*d*_*i*_|). Positive values indicate deviation above the reference line and negative values deviation below it.

### Paddle–ball coupling

For the coupling analysis, ball and paddle trajectories were resampled onto a uniformly spaced temporal grid (0.025 s resolution, 40Hz) using linear interpolation. This ensured constant temporal spacing required for smoothing and enabled comparison across trials. Trajectories were then smoothed using a Savitzky– Golay filter (polynomial order *p* = 3, window length *n* = 11 samples) to reduce noise while preserving movement dynamics.

For each trial, the horizontal position at which the ball trajectory intersected the initial vertical position of the paddle was estimated by interpolation:

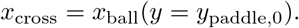

Trajectories were subsequently normalized by (i) horizontally flipping trials to a common reference frame, (ii) centering positions relative to their initial values, and (iii) scaling by the crossing position *x*_cross_. After normalization, positions are expressed as dimensionless displacements, where 1 corresponds to the horizontal distance to the crossing point.

The coupling index was defined as:

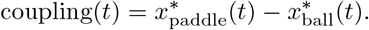

Let *A*^+^ and *A*^*−*^ denote the positive and negative areas under the coupling curve, computed as:

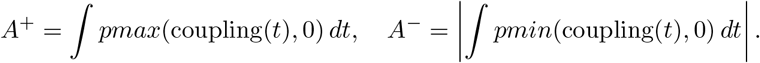

The coupling ratio (*AUC*) was defined as:

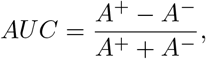

which is bounded in [*−*1, 1]. Values close to +1 indicate that the paddle predominantly leads the ball trajectory, values close to *−*1 indicate that it lags behind, and values near 0 indicate balanced coordination.

## Appendix C: Statistical models and supplementary analyses

This appendix reports full model specifications, parameter estimates, pairwise comparisons, and supplementary figures complementing the main results presented in the manuscript.

## Supplementary figures

**Fig. 2.**
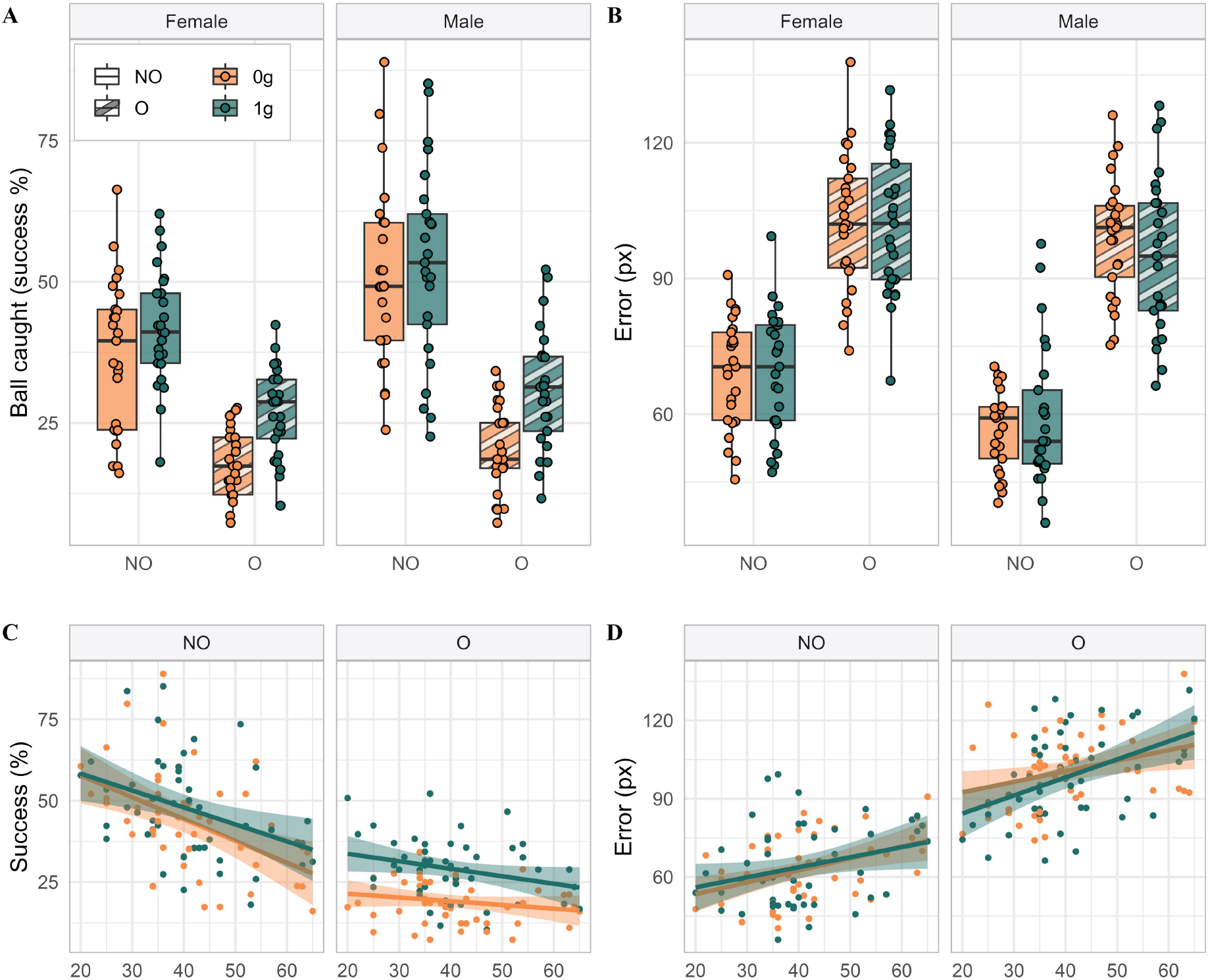
Supplementary analyses of success rate and interception error as a function of gender. Panel A shows the percentage of successful interceptions (ball caught) as a function of occlusion (NO, non-occluded; O, occluded) and gravity (0g, orange; 1g, green), shown separately for female and male participants. Panel B shows the interception error (in pixels) for missed trials only, under the same conditions and conventions. Boxplots represent the distribution of participant-level data with individual observations overlaid. Panels C and D show success rate and interception error, respectively, as a function of age (x-axis) and occlusion condition, separately for each gravity level; regression lines with 95% confidence intervals are shown for descriptive purposes. These analyses were conducted as exploratory supplementary analyses and are not discussed in the main text; no inferential statistics are reported.

**Fig. 3.**
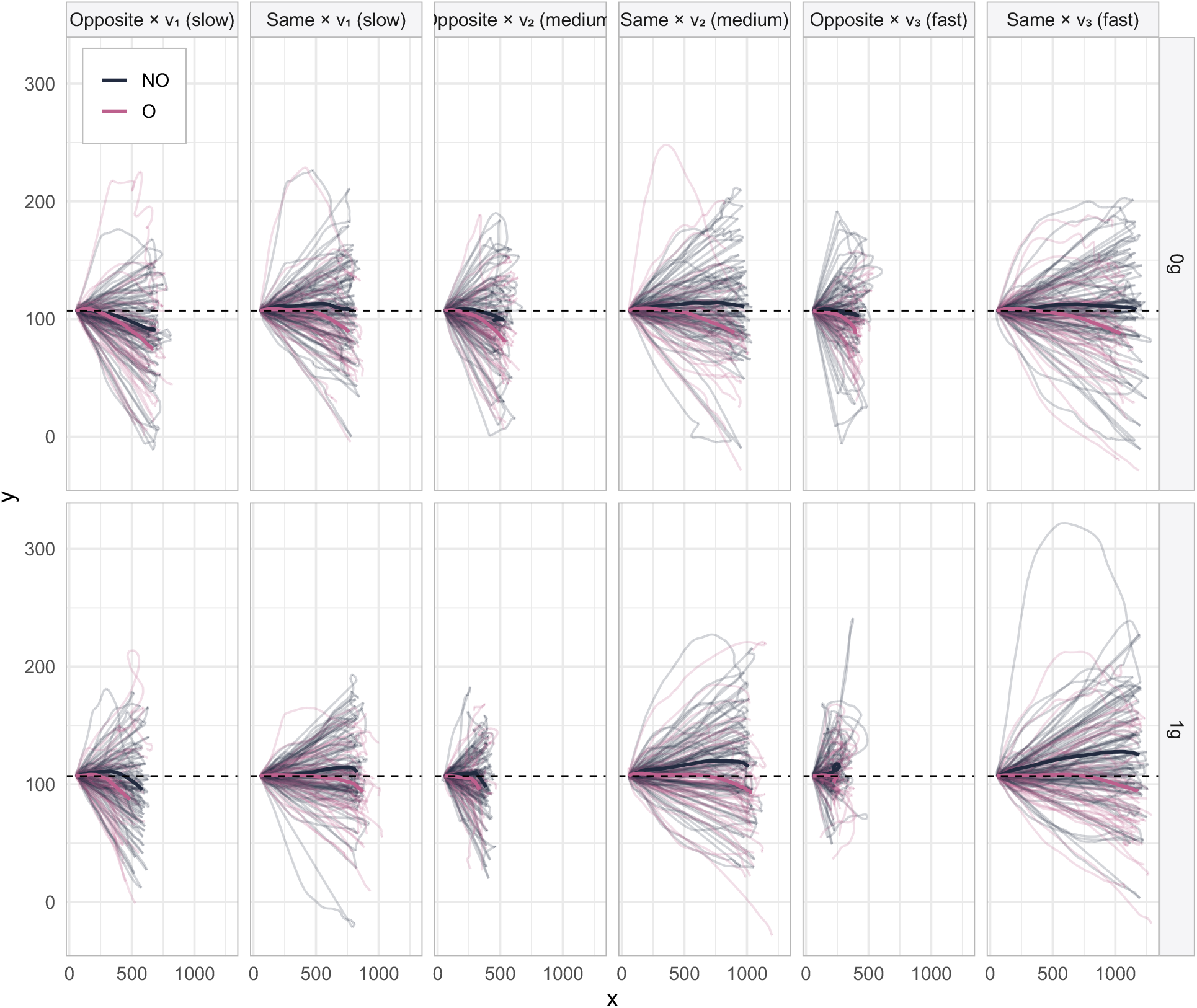
Supplementary visualization of raw and average cursor trajectories across experimental conditions. Each panel shows the full set of individual mouse trajectories (light grey lines) alongside the condition average (dark line) in the two-dimensional display space (*x* = horizontal position in pixels; *y* = vertical position in pixels).

